# DNA methylation reveals distinct cells of origin for pancreatic neuroendocrine carcinomas (PanNECs) and pancreatic neuroendocrine tumors (PanNETs)

**DOI:** 10.1101/2020.06.12.146811

**Authors:** Tincy Simon, Pamela Riemer, Katharina Detjen, Annunziata Di Domenico, Felix Bormann, Andrea Menne, Slim Khouja, Nanna Monjé, Liam H. Childs, Dido Lenze, Ulf Leser, Armin Jarosch, Florian Rossner, Markus Morkel, Nils Blüthgen, Marianne Pavel, David Horst, David Capper, Ilaria Marinoni, Aurel Perren, Soulafa Mamlouk, Christine Sers

## Abstract

Pancreatic Neuroendocrine Carcinomas (PanNECs) are high-grade, poorly-differentiated tumors grouped together with Pancreatic Neuroendocrine Tumors (PanNETs) and placed within the Pancreatic Neuroendocrine Neoplasms (PanNENs) WHO tumor classification. Despite recent studies suggesting the endocrine origin of low-grade PanNETs, high-grade PanNEC origin remains unknown. DNA methylation analysis using the Illumina 850K beadchip array was conducted on 57 PanNEN samples, including 14 PanNECs. Distinct methylation profiles separated PanNEN samples into two major groups, clearly distinguishing high-grade PanNECs from other PanNETs including high-grade NETG3. DNA mutations, copy number changes and Immunohistochemistry of pancreatic cell-type markers PDX1, ARX and SOX9 were utilized to further characterize PanNECs and their hierarchical cell of origin in the pancreas. Phylo-epigenetic and cell-type signature features using methylation data from normal alpha, beta, acinar and ductal adult cells indicate an exocrine cell of origin for PanNECs, thus separating them in cell lineage from other PanNENs of endocrine origin. Our study provides a robust and clinically relevant method relying on methylation profiles to clearly distinguish PanNECs from PanNETG3s to improve patient stratification and treatment.

## Introduction

Pancreatic Neuroendocrine Neoplasms (PanNENs) have undergone several classification changes according to consensus guidelines ^1–3^. In 2017 these rare tumors were grouped by the WHO, with respect to proliferation index and morphology, into well-differentiated Pancreatic Neuroendocrine Tumors (PanNETs) and poorly differentiated Pancreatic Neuroendocrine Carcinomas (PanNECs)^4^. PanNETs were further divided into G1, G2, or G3 tumors, with an increasingly malignant nature depending on their proliferation index (Ki67 <3%, 3-20% and > 20% respectively)^4,5,6^. PanNECs were all G3 due to a high proliferation rate (Ki67 > 20%) combined with poor differentiation of the cells, resulting in an aggressive phenotype and poor prognosis ^7^. Thus, the new classification introduced the important biological distinction between PanNETs of grade G3 (NETG3s) and PanNECs, despite their similar proliferation ^8^. Histologically, however, the distinction between G3 PanNETs and PanNECs remains difficult and ambiguous in a high number of cases, leading to misclassification^9^.

All PanNENs are malignant tumors and pancreatic in origin, however the field is rapidly acknowledging a clear distinction between poorly differentiated PanNECs and well-differentiated PanNETs ^2^. Compared to PanNETs, PanNECs are highly aggressive and rapidly fatal, and most patients die within one year of diagnosis ^10,11^. PanNECs are also more responsive to platinum-based treatments than PanNETs, at least initially ^12,13^. Unlike PanNETs, which carry mutations in *MEN1, ATRX, DAXX*, PanNECs have a mutational profile similar to Pancreatic ductal adenocarcinoma (PDACs), characterized by genomic alterations in *KRAS, SMAD4* and *TP53.* They additionally display loss of Rb1, further distinguishing them from PanNETs ^14^. PDACs have been shown to originate from normal ductal or acinar cells ^15,16^, while G1 and G2 PanNETs have been shown to originate from endocrine cells of the pancreas ^17,18,19^. As yet, no study has attempted to identify the origin of PanNECs.

The pancreas consists of two compartments: an endocrine compartment (islets of Langerhans) with alpha (α), beta (β), delta (δ), epsilon (ε) and polypeptide producing (PP) cells ^20–23^ which produce glucagon, insulin, somatostatin and pancreatic polypeptide ^24^ respectively, and an exocrine compartment composed of ductal, centro-acinar and acinar cells ^20–23,25^. Many transcription factors are expressed and repressed in a spatio-temporal manner in order to form and maintain adult pancreatic cell types (reviewed by Cano et. al.^26^ and Arda et. al. ^27^). The transcription factor PDX1 is critical for maintaining β cells ^28–33,34^ while ARX, upstream of IRX2, is a well-established α-cell-specific transcription factor ^28–30,35^ . Presence of NKX6-1, NKX2-2 and PAX6 further maintains the endocrine lineage for α and β cells ^36,37^. SOX9, a downstream target of NOTCH, is required for the establishment of cell fates of endocrine and exocrine cells ^38,39^. While SOX9 is absent in the committed endocrine precursors ^40^, it is an important player in pancreatic ductal and centro-acinar cell development but not found in adult acinar cells ^25,41^. Importantly, Kopp et al. have shown a relationship between oncogenic *KRAS* and *SOX9* expression in the formation of premalignant PDAC lesions ^42^. The sub-classification of PanNENs routinely relies on proliferation, morphological markers, immune phenotype, and symptoms associated with excessive hormone secretion. However, as most high-grade PanNETs and indeed all PanNECs are non-functional tumors, a more robust method for their distinction is required ^43^.

In this study we utilized genetic and epigenetic profiling, including Illumina 850K beadchip array methylation profile analysis, to classify PanNECs in relation to PanNETs of all grades and to identify the possible cell of origin of PanNECs. We find that PanNECs have distinct DNA methylation profiles from PanNETs, providing a means to distinguish histologically similar PanNECs and high-grade PanNETs. Furthermore, similarities between acinar cell and PanNEC methylation profiles point to an exocrine cell origin for this subtype of pancreatic carcinomas.

## Results

### Sample characteristics

To characterize and classify PanNEN tumors we performed methylation analysis using the Infinium Methylation EPIC (850K) beadchip platform in addition to DNA high-depth panel sequencing of 57 PanNEN. The cohort consisted of 43 PanNETs including NETG1 (n = 18), NETG2 (n = 13), NETG3 (n = 12), and PanNECs (n = 14). Forty one samples were collected from primary PanNENs and 16 from metastases (12 liver, 2 lymph nodes, 1 bladder and 1 peritoneal metastasis) (representative H&E sections in Fig. S1a). For two patients we had two separate samples; PNET77 (primary and liver metastasis) and PNET56 (two liver metastases excised one year apart). As healthy controls, we used either matched adjacent healthy (15), distant healthy tissue (29) or blood (3) (Fig. S1b). Mean tumor cell content, evaluated by a pathologist, was 85.4% +/- 11%. All samples came from formalin-fixed paraffin-embedded tissue. Samples used for each assay and their characteristics are depicted in Fig. 1a and Table 1. Further details can be found in Supplementary Table 1.

**Fig. 1.**
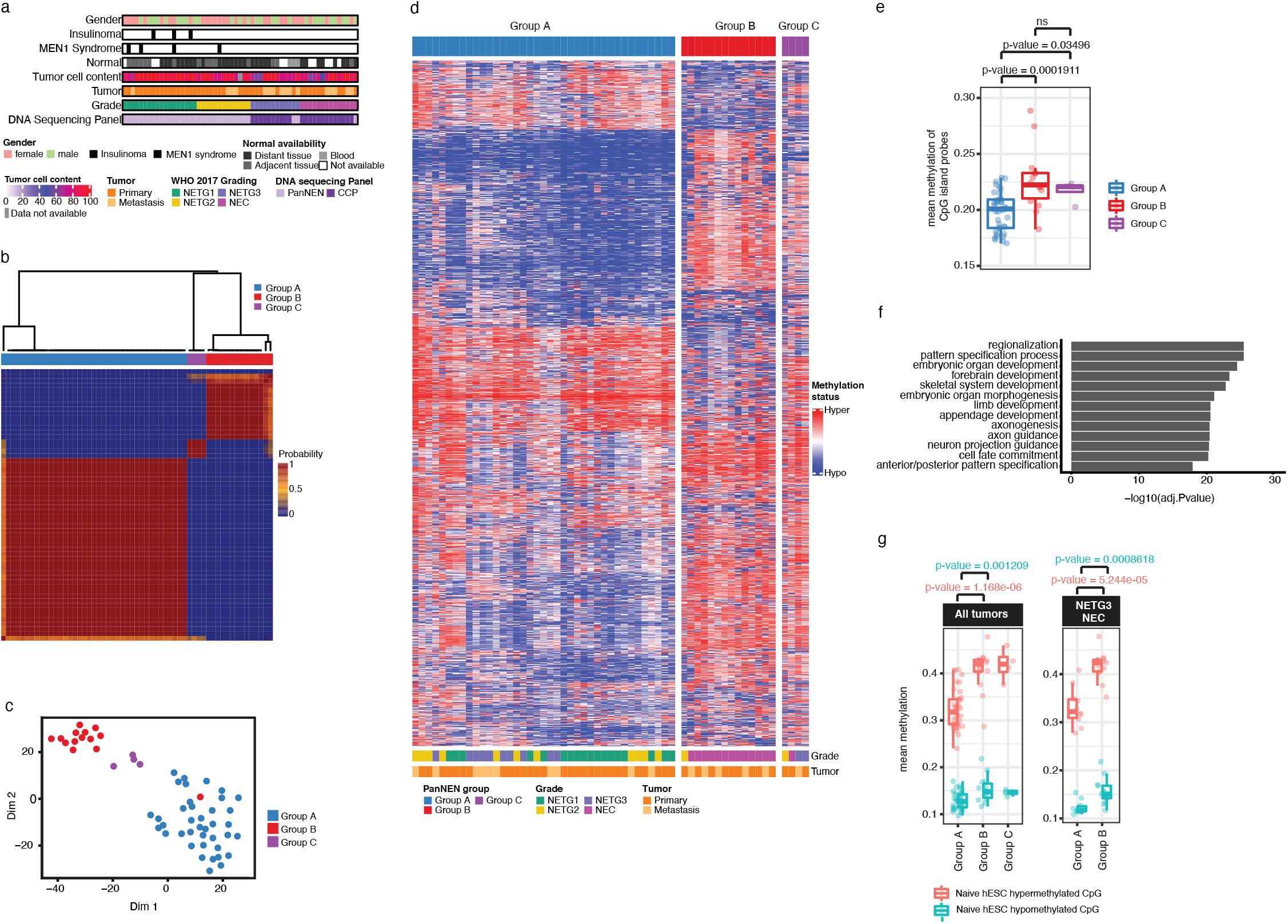
PanNEN tumors subdivide into two main methylation groups. **a**. Characterization of PanNEN cohort. Normal: matched healthy tissue, tumor cell content: tumor cell fraction in the material used for analysis as estimated by a pathologist. **b.** Unsupervised class discovery using 10000 (10K) most variable methylation probes. Heatmap displays pairwise consensus values of the samples. **c.** tSNE representation of PanNEN subgroups using 10K most variable probes. Subgroup annotation is used as colors for each sample (dot). **d.** Heatmap displaying the methylation status of 10K variable probes in each of the Groups; A, B and C. Methylation beta value was used to perform hierarchical clustering separately on each subgroup, identifying closely similar samples. Hierarchical clustering was performed on the probes; color range blue to red represents methylation beta value, columns indicate samples and rows methylation probes. **e.** Mean methylation of CpG island probes in PanNEN subgroups. Boxplot represents distribution of mean methylation of CpG island associated probes. Each dot depicts the mean value of CpG island associated probes in a given sample. **f.** GO ontology analysis of 10K most variable probes, representing top 12 GO terms based on -Log10P-value. **g.** Mean methylation of hESC associated hypermethylated and hypomethylated probes in PanNEN subgroups. Boxplot represents distribution of hypermethylated (red) and hypomethylated CpG probes of hESC (blue) in all samples (left panel), and only PanNETG3/ PanNEC samples from Group A and B respectively (right panel); Each dot depicts the mean value of probes in each hESC category for a given sample. Statistical analysis was performed on the difference in distribution between subgroups using a two-sample Wilcoxon test. Box shows 25th and 75th percentiles and sample median as horizontal line, whiskers show maximum and minimum value that is 1.5 times the interquartile range over the 75th and 25th percentile, respectively.

**Table 1.** Cohort characteristics

| Characteristics | Variable | count | (%) |
| --- | --- | --- | --- |
| Gender | Female | 27 | 47.4 |
|  | Male | 20 | 35.1 |
|  | Unknown | 10 | 17.5 |
| Normal | Normal adjacent tissue | 15 | 26.3 |
|  | Normal distant tissue | 29 | 50.9 |
|  | No normal | 10 | 17.5 |
|  | Peripheral blood | 3 | 5.3 |
| Tumor | Metastasis | 16 | 28.1 |
|  | Primary | 40 | 70.2 |
|  | Unknown | 1 | 1.8 |
| Location | Head | 17 | 29.8 |
|  | Tail | 14 | 24.6 |
|  | Liver | 12 | 21.1 |
|  | Body | 2 | 3.5 |
|  | Lymph node | 2 | 3.5 |
|  | Multiple primary lesion location | 2 | 3.5 |
|  | (Other) | 8 | 14.0 |
| Grade | NETG1 | 18 | 31.6 |
|  | NETG2 | 13 | 22.8 |
|  | NETG3 | 12 | 21.1 |
|  | NEC | 14 | 24.6 |
| Diagnosis | Insulinoma | 3 | 5.3 |
|  | MEN1 Syndrome | 4 | 7.0 |

### DNA methylation classification identifies a distinct PanNEC subgroup within PanNEN samples

To determine PanNEN subtypes, we performed unsupervised class discovery using the 10,000 (10K) most variable probes and defined three distinct groups at the methylation level; Groups A, B and C (Fig. 1b; Fig. S1c and methods for details). A t-distributed stochastic neighbor embedding (tSNE) analysis showed consistent segregation of the samples, affirming the presence of distinct methylation patterns in the groups identified (Fig. 1c). The methylation beta values from the 10K most variable probes revealed notable distinction between the groups (Fig. 1d), as Group A was composed of 39 well-differentiated PanNETs of all grades including 10 NETG3, while Group B harbored 13 from a total of 14 PanNECs and one NETG2 sample. Group C consisted of only four samples; two NETG3s, one NETG2, and one NEC (supplementary table 1 for details). Patient survival data confirms the distinct separation of Group A and B enriched in PanNET and PanNEC, respectively (Patient survival graphs according to tumor grade and methylation group in Fig. S1d). Mean CpG island methylation was significantly higher in Group B and Group C compared to Group A (Two-sample Wilcoxon Test: p-value = 0.0001911 between Group A and B and p = 0.03496 between Group A and C) (Fig. 1e). Importantly, the methylation profiles separate high-grade NETG3 from NEC tumors as they were assigned with high precision to Group A and Group B, respectively.

Gene Ontology (GO) analysis of the genes associated with the 10K probes significantly enriched for terms involved in organ development and specifically, neurogenesis associated biological processes (Supplementary table 2; Fig. 1f). Recent work evaluating the transition of primed human Embryonic Stem Cells (hESC) to naive state demonstrated a gradual and stable acquisition of CpG hypermethylation in genes associated with development which was mirrored in multiple cancer entities ^44^. We found that the naive hESC-associated and hypermethylated CpGs were significantly more hypermethylated in Group B compared to Group A tumors (Fig. 1g left panel). A closer analysis of NETG3 samples in Group A compared to NEC samples from Group B maintained a similar significant difference in the distribution of hypermethylated CpGs of naive hESC (Fig. 1g right panel), highlighting the difference in developmental states between these histologically similar high-grade PanNEN tumor types.

Taken together, we have identified three methylation groups, reflecting the histopathologically distinct PanNETs and PanNECs, with PanNECs in Group B mirroring the epigenetic signature associated with naive hESCs.

### Distinct recurrent mutations separate PanNENs in Groups A and B

To further characterize the PanNEN samples at the mutational level, we employed high-depth panel sequencing. We utilized the commercially available Comprehensive Cancer Panel (CCP), and a custom PanNEN panel covering 47 additional genes known to be relevant in PanNEN biology (Fig. S2a, supplementary table 3, details in methods). Together, the panels covered 432 genes.

In total, 50 genes were found altered in 41 of the 57 PanNENs of our cohort. Classical alterations associated with PanNETs were enriched in Group A, as recurring mutations in *MEN1, DAXX, ATRX* and *TSC2* were found in 13, 6, 4 and 4 of the 39 Group A samples, respectively. In contrast, mutations in *DAXX* and *ATRX* were absent in Group B, while only one Group B sample each was mutated in *MEN1* and *TSC2* (Fig. 2a shows recurring mutations; detailed view of all mutations in Fig. S2b and supplementary table 4). In addition, four Group A samples contained aberrations in *VHL* and two samples contained *PTEN* mutations. In contrast, *KRAS* (5 out of 14 samples, comprising G12D, G12R, G12V and Q61D mutations), *SMAD4* (2) *and TP53* (3) mutations were exclusively seen in Group B. In total, 16 samples harbored no known driver mutations detected by our panel, and this included the single non-PanNEC sample in Group B (note that all mutations discussed were either non-synonymous, deletions or indels, for allelic frequencies see Fig. S2c). Two patients in our cohort had multiple samples: PNET77 and PNET56. Patient PNET77 had tissues of primary tumor and liver metastasis (PNET77P and PNET77M, both samples are NETG3 in Group C) surgically removed two years apart. Neither sample displayed mutations picked up by our panel. In contrast, Patient PNET56 had two liver metastasis samples in our cohort; PNET56P1 and P2 (both NETG3s in Group A) removed one year apart. Both samples carried the same alterations in TSC2 and BRD3. Collectively, the mutational profiles uncovered key molecular distinctions between the Group A and B samples, which were enriched for aberrations in *MEN1, DAXX* and *ATRX* and *KRAS*, *TP53* and *SMAD4*, respectively.

**Fig. 2.**
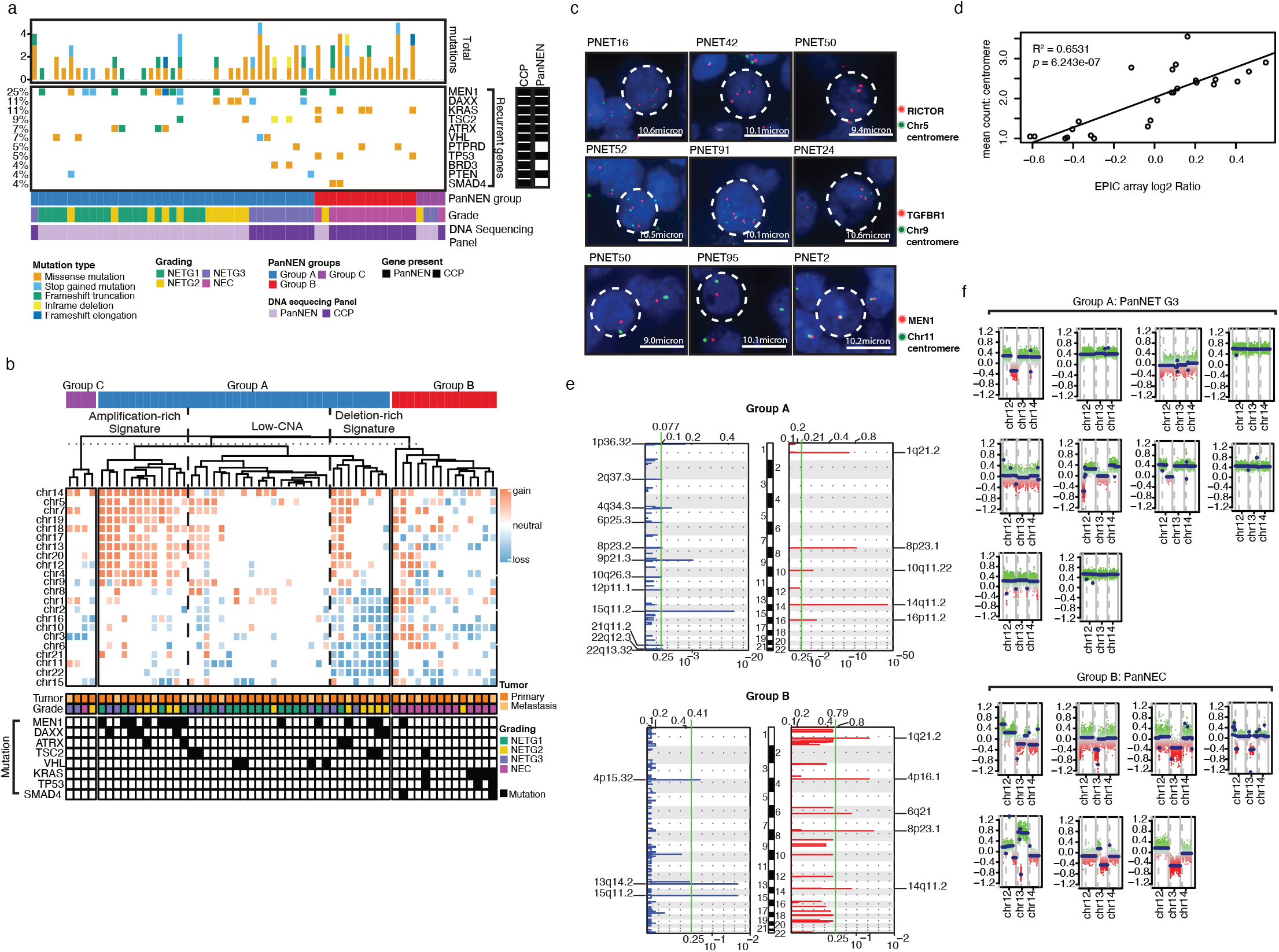
Genetic aberrations distinguish Group A and Group B. **a**. Recurrent gene mutations in PanNEN subgroups. Barplot depicts the mutational frequency in a given sample, colored according to variant type (top panel). Percentage value per row denotes the frequency at which the respective gene is aberrated in the cohort (left); recurrent mutation variants of each patient, colored according to variant type (middle panel). White spacing: no mutation identified in the regions of the targeted gene; row annotations indicating which sequencing panel was used:PanNEN panel, CCP panel, or both (right); PanNEN subgroup, tumor grade and gene panel used for mutational analysis for each sample. **b**. Whole chromosomal aberrations in PanNEN subgroups. Hierarchical clustering of mean log2 ratios of chromosomal segments per autosome is displayed in a heatmap; dotted line represents cut off used to identify amplification, low-CNA and deletion-rich signatures; column annotation (bottom) are tumor grade, tumor type and recurrently aberrated genes. **c**. Representative images of FISH validation of whole chromosomal aberrations in PanNEN subgroups. Green: centromere of chr5 (top panel), chr9 (middle panel) and chr11 (bottom panel). **d**. Correlation of log2 ratio and FISH count. Linear regression of mean copy number count of centromere derived from FISH (y-axis) and mean log2 ratios of chromosomal segments per autosome (x-axis); diagonal line: best fit model for the data. **e**. Significant focal copy number aberrations in PanNEN subgroup. Focal aberrations in Group A (top panel) and Group B (bottom panel) displayed; Blue: focal copy number losses, red: gain. Log2 ratio range at the top and q-value at the bottom. Green: q-value cut off at 0.25 to call significant. Significantly aberrated focal regions are indicated (left and right). **f.** Chromosome 12, 13, and 14 copy number status in NETG3 (top panel) and NEC (bottom panel). Intensity values of each bin are plotted in colored dots; each color indicates ‘methylated’ and ‘unmethylated’ channels of each CpG; Segments are shown as horizontal blue lines.

In addition to the genetic differences detected between Group A and Group B PanNENs, we also found mutations shared between the groups. Common alterations were found in genes involved in chromatin-remodeling processes including mutations in *EP300, BRD3, AFF1, MLLT10, PSIP1, SETD2* and *KMT2C,* which were represented in both Group A and Group B. Three alterations targeting PI3K subunits such as *PIK3C2B* and *PIK3CG* were found in both groups (Fig. S2b). These mutations point to common mechanisms driving PanNEN development irrespective of the differences between Group A and Group B tumors detailed above.

### Copy number alterations separate PanNECs from PanNETs

To explore copy number alterations (CNAs) in PanNENs, we first inferred methylation log2 ratios from signal intensities. For analysis of whole chromosomal copy number changes within each group, we calculated the average log2 ratio of CNA segments per chromosome and subsequently performed unsupervised clustering. Within Group A we found three profiles: amplification-rich, deletion-rich and low-CNA signature (Fig. 2b). The amplification-rich signature predominantly harbored copy number gains in chromosomes 14, 5, 7,19, 18, 17, 13, 20, 12, 4 and 9. Samples in the deletion-rich signature carried recurrent deletions of chromosomes 6, 1, 22, 8, 1, 2, 16, 10, 3, and 21, as also validated by fluorescence *in-situ* hybridization (FISH; Fig. 2c, Fig. S3a and b). Mean signal counts per sample from FISH and mean log2 ratio from CNA analysis were correlated and showed a regression coefficient of R^2^ =0.6531 and p=6.243 × 10^-7^ (Fig. 2d). Recurrent mutations of tumor suppressor genes *MEN1, DAXX, TSC2,* and *VHL* were enriched in amplification-rich and deletion-rich signature (Fig. 2b, lower column annotation). The low-CNA signature contained few aberrations in most chromosomes with no clear recurrences, and this signature was a predominant feature of NETG1 tumors of Group A. Tumors in Group B and Group C were defined by few recurrent whole chromosomal aberrations, likely due to the low number of samples in the groups.

In addition to whole chromosomal aberrations, we investigated focal CNA using GISTIC (Fig. 2e). Chromosome regional gains of 1q21.2, 8p23.1 and 14q11.2, and deletion of chromosomal region 15q11.2 were significantly associated with both Group A and Group B (supplementary tables 5 and 6). Group A had unique significant gains in chromosomal regions 10q11.22 and 16p11.2, which were not present in Group B. In contrast, only Group B showed focal gains in 4p16.1 and 6q21. Chromosomal deletions exclusive to Group A were in regions 1p36.32, 2q37.3, 4q34.3, 6p25.3, 8p23.2, 9p21.3, 10q26.3, 12p11.1, 21q11.2, 22q12.3 and 22q13.32, whereas Group B carried focal deletions in 4p15.32 and 13q14.2. Deletion of 9p21.3 in Group A resulted in loss of *CDKN1B and CDKN2A*, two key regulators of the cell cycle (supplementary table 5). Interestingly, deletion of 13q14.2 affecting *RB1,* also an important cell cycle regulator, was seen in 50% of Group B. (Fig. 2f, supplementary table 6). On closer examination, we found that NETG3s in Group A did not show losses of *RB1* region (Fig. 2f lower panel). Therefore, while Group A enriched for recurrent whole chromosomal and focal copy number aberrations, Group B specifically showed significant focal aberrations and exclusively harbored *RB1* loss associated with PanNEC tumors.

### PanNETs in Group A harbor endocrine cell of origin signatures

Cell-of-origin studies in PanNEN have been thus far restricted to well-differentiated PanNETs. Therefore, we investigated cell-of-origin methylation patterns in our diverse PanNEN tumors. Our first approach explored established markers of pancreatic cell lineages. We curated a list of 174 markers from PanglaoDB ^37^ uniquely expressed in each differentiated cell type of the adult pancreas (marker list in supplementary table 7; see method for details). We called differentially methylated probes (DMPs) from our samples (supplementary table 8) and identified genes overlapping with the curated list. In total, we detected 122 markers associated with the 770 DMPs from our samples. Among the identified probes, we determined genes which showed a significant difference between Group A and Group B (-log_10_P > 5 and Δbeta > |0.25|) (Fig. 3a, Fig.S4a, supplementary table 9). We identified 85 significantly enriched probes (red points in Fig. 3a) associated with 23 markers of α, β, ɣ, δ, ductal, acinar and Islet Schwann cells (Fig. S4b). Among these, α cell markers such as *IRX2, TTR* and *GLS* were hypomethylated across Group A samples (Fig. 3b). Multiple probes associated with *IRX2* showed the most significant difference between Groups A and B (Fig. 3a). Indeed, Group A tumors were consistently hypomethylated in promoter associated probes of *IRX2*, while Group B showed strong hypermethylation (Fig. 3b) Although *PDX1* was not differentially methylated between Group A and Group B, the 10K probes establishing the groups showed variable methylation patterns of this gene within the groups (Fig. 3c and Fig. S4c for sample IDs). We saw that Group A tumors separated with respect to *IRX2* and *PDX1* methylation status, whereby one subgroup carried hypomethylation of both *IRX2* and *PDX1*, while in the second subgroup *PDX1* was hypermethylated. The latter subgroup was enriched for *MEN1, DAXX* and *ATRX* mutations, while recurring *VHL* mutations were seen in the prior subgroup (Fig. 3c). In agreement with these data, and with previous studies ^45^, the transcription factor ARX was also found expressed in a subgroup of Group A tumors and ARX+PDX1- phenotype was significantly more common compared to Group B (p-value=0.02036: Fisher’s exact test) (Fig. 3d and Fig. S4e and S4f). In addition, genes associated with endocrine cell lineage maintenance, such as *PAX6, NKX6-1* and *NKX 2-2* were mainly hypomethylated in Group A, except for NETG3 samples (PNET42, PNET57, PNET61, PNET56P1, PNET56P2 and PNET107), and one NETG2 (PNET24), which showed hypermethylation of *PAX6*, *NKX6-1* and *NKX2-2* genes similar to Group B samples (Fig. 3b). In contrast, Group B, comprising almost exclusively of PanNECs, was characterized by hypermethylation of all DMPs under investigation, with the exception of *KRT7*.

**Fig. 3.**
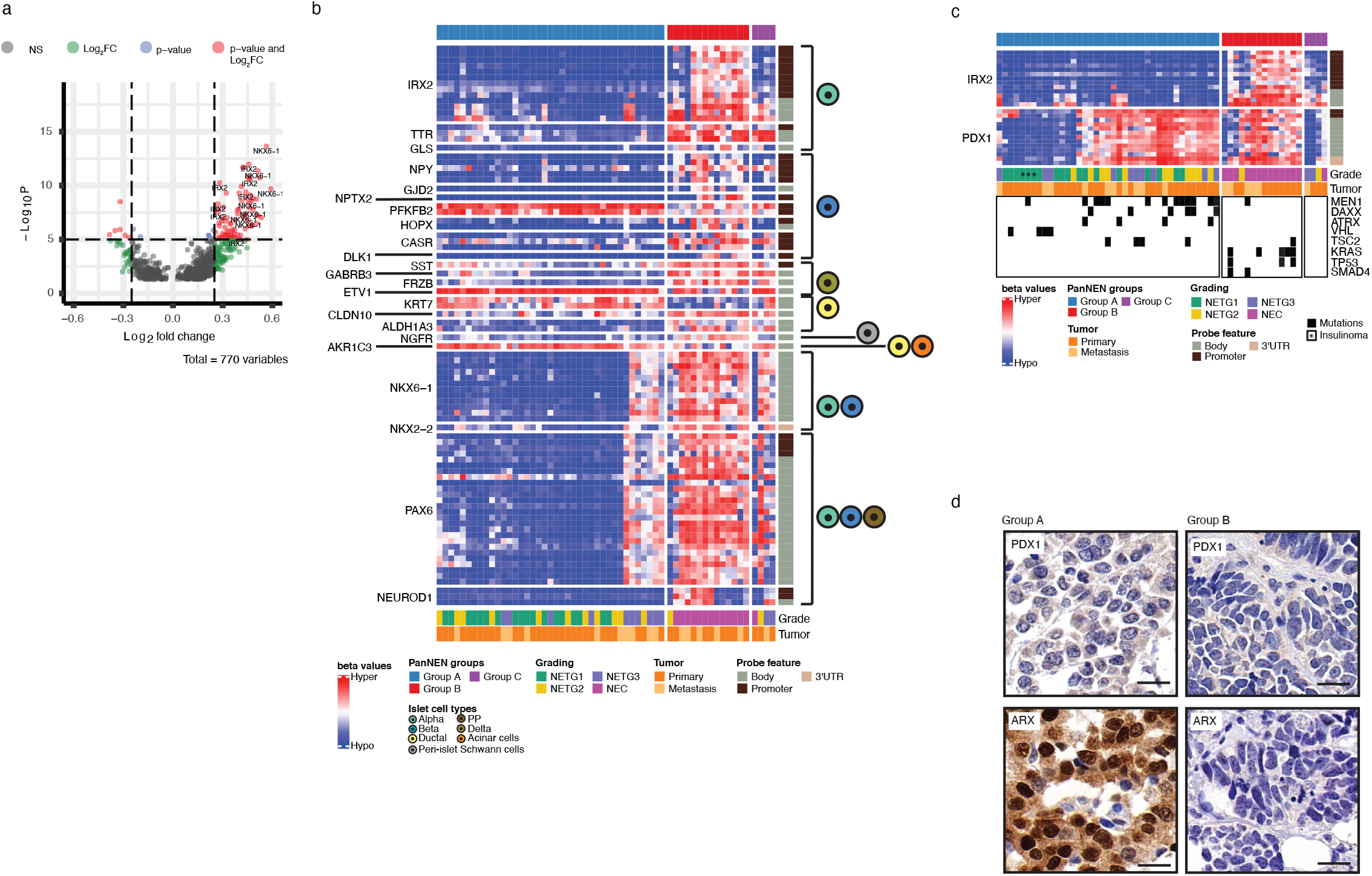
Cell marker analysis in the PanNEN subgroup identifies endocrine features in Group A. **a**. Differentially Methylated Probes (DMPs) from our cohort associated with pancreatic cell markers (n=770). Dots: intersect between -Log10 P value and the log2 fold change (FC) for a given probe. Cut off for significant -Log10 P-value: 5 (adjusted p-value: 10^-6^); cut-off for significant log2FC: >|0.25|. Red: probes passing both cut-offs; Green: probes only surpassing the log2FC threshold; Blue: probes that only have a significant p-value; Grey: probes that did not pass either of the cut-offs; Significantly associated DMP probes of IRX2 and NKX6-1 labeled in the plot. **b.** Methylation beta value of significant DMPs of pancreatic cell markers of PanNEN subgroups. Heatmap displaying the methylation beta values of DMPs (row) in each sample (column). The heatmap color represents the methylation beta value; row annotation identifies the genomic region associated with the probe (panel on the right) in addition to which cell type the gene shows specificity in gene expression (according to PanglaoDB). **c.** Methylation beta value of probes associated with *IRX2* and *PDX1* in PanNEN subgroups. DMP probes of *IRX2* and 10K probes associated with *PDX1* (rows) for each sample (column); rows are split at each associated gene (left); columns are split at each subgroup (bottom); row annotation: genomic region associated with the probe (panel on the right); column annotation (bottom): the tumor grade, tumor type as well as the recurrently aberrated genes in the context of *IRX2* and *PDX1* methylation state. **d.** Representative IHC of *ARX* and *PDX1* in PanNEN subgroups. Scale bar: 20µm.

### PanNECs in Group B display an acinar-like cell signature and differ from PDACs in ductal-like cell signatures

In order to identify the cell of origin for samples from Group B, we next extended our analysis to include normal pancreatic methylation profiles. We obtained Illumina 450K array methylation profiles of presorted normal pancreatic cell types: α (n =2), β (n =3), acinar (n =3) and ductal (n =3) cells ^46, 47^. We identified 46,500 DMPs differentiating the cell types from one another (adjusted P value < 0.01, absolut Δ beta > 0.2, supplementary tables 10 to 15). We determined Pearson distance between the tumor samples and the pancreatic cell types using the DMPs, and constructed a phylo-epigenetic tree (Fig. 4a). Two main branches separate the whole cohort; the lower branch consists completely of Group A, and clustered closely with β and α cells. The insulinomas grouped together and maintained the closest distance to the normal endocrine cell type. The second main branch of the phylogeny consists of all three Groups, with NETG3 samples mainly clustering together. Strikingly, all tumors from Group B, except PNET58, distinctly clustered with ductal and acinar cells, forming a separate clade together, indicating a clear resemblance to the exocrine cells of the pancreas and evidently distant from the endocrine cells of the pancreas. Interestingly, PNET58 was ARX+ and PDX1+ (Fig. S4d).

**Fig. 4.**
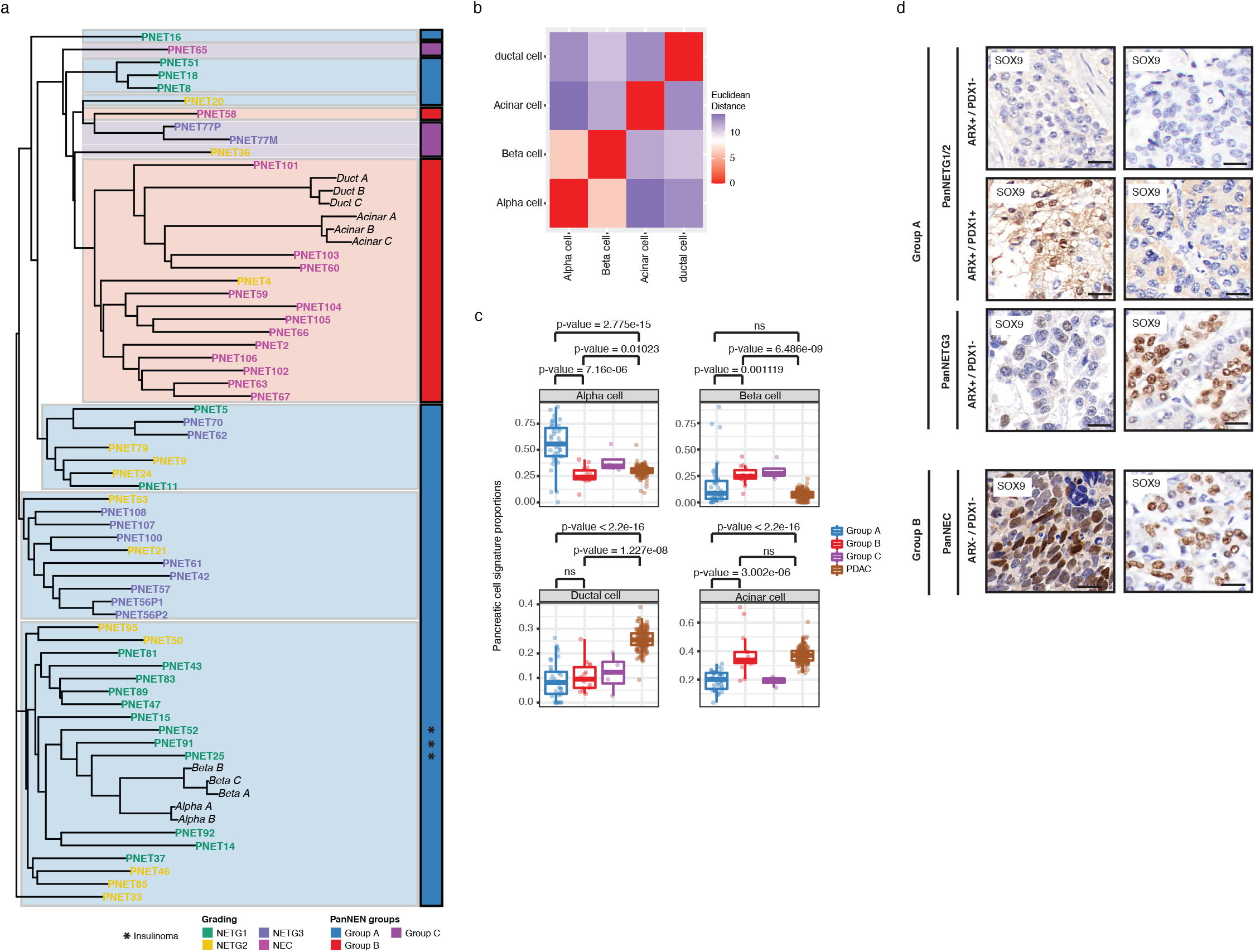
Cell-of-origin analysis using normal cell type methylation profiles and SOX9 in the PanNEN subgroup indicate exocrine lineage for Group B tumors. **a.** Phylo-epigenetic analysis of PanNEN tumors and normal pancreatic cell types. Pearson distance between the samples computed using the differentially methylated CpGs between normal α-, β-, ductal and acinar cells samples (*n* = 46,500, adj. *p* value < 0.01 and |Δβ | > 0.2) and neighbor-joining tree estimation was performed to determine clades and branching. Bottom annotation identifies the PanNEN subgroups and highlights the main branch from which the closely associated samples of a given subgroup split. **b.** Euclidean distance between each cell type was computed and correlation matrix of the distances are displayed here. The heatmap color depicts the distance between a given normal cell-type pair. **c**. Boxplot representing distribution of the proportion of atlas signature of α-, β-, ductal and acinar cells (each main box) in the subgroups and PDACs; each dot depicts the proportion of atlas signature of the respective cell type in a given sample; statistical analysis was performed on the difference in distribution between subgroups using a two-sample Wilcoxon test. **d.** IHC of SOX9 in PanNEN subgroups. Representative images for each subgroup; scale bar: 20µm.

We next applied an independent method to determine the normal cell signature composition of our PanNEN samples. As a reference, we utilized 6096 CpG probes defined by Moss et al. which distinguish ductal, acinar and β cells ^48^. For the missing α cell reference we utilized the 450K profiles mentioned in the previous section ^46^ and determined the methylation values for the CpGs which differentiated β, acinar and ductal cells (Supplementary table 16). Using euclidean distance analysis, we confirmed that the reference cell types could be clearly discerned from one another (Fig. 4b). Given the reference and the tumor samples, we ran a deconvolution algorithm which models the profile of the tumors as a linear combination of the methylation profiles of the cell types (details in methods). Under this assumption, the method determines the methylation signature proportion using non-negative least squares linear regression (NNLS). In addition to the grouped samples from our PanNEN cohort, we used 167 PDACs as an independent pancreatic tumor type ^49^. Group A PanNETs displayed significant enrichment of α-cell signature composition compared to Group B and to a lesser extent to Group C (Fig. 4c and Fig. S5a and Fig. S5b). Group B NECs, on the other hand, showed a significant increase in acinar cell signature proportion when compared to Group A and to Group C. Interestingly, the acinar cell composition of Group B was similar to that found in PDAC tumors. However, the ductal signature composition of PDAC tumors was significantly higher than other groups of PanNENs, distinguishing this pancreatic exocrine cancer from PanNENs and specifically from Group B. NETG3 samples from Group A contained similar profiles to other PanNET samples, and showed no resemblance to the acinar cell similarity of PanNECs (Fig. S5c).

Using the same method, we went back to look at the two subgroups within Group A which showed differences in *PDX1* methylation patterns (Fig. 3c). We found an α-like cell profile signature more significantly enriched in samples with hypermethylation of *PDX1*, and a β-like and intermediate cell profile in samples with hypomethylation of both *PDX1* and *ARX* (Fig. S5d).

### Group B PanNECs display SOX9 patterns similar to exocrine cells

The pattern of Group B acinar similarity to PDACs compelled us to investigate where this similarity originated from. An earlier study linked SOX9 and formation of precursor lesions of PDACs from acinar cells together with expression of oncogenic KRAS ^42^. To identify whether the PanNECs in Group B display SOX9, we performed IHC for representative samples of the cohort. From 11 PanNEC samples of Group B, 9 samples were positive for SOX9 staining (representative images in Fig. 4d, Fig. S4e). In Group A, we found 4 /17 samples (three NETG3s and one NETG2 (an insulinoma and MEN1 syndrome tumor)) staining for SOX9, albeit with a heterogeneous and low intensity staining, unlike the strong homogenous expression found in Group B positive tumors (representative images in Fig. 4d and Fig. S4f). To further link the endocrine/exocrine feature with regard to *SOX9* expression, we additionally analyzed IHC of ARX and PDX1 on the same samples. The Group A SOX9+ NETG3 were additionally ARX+. As for Group B SOX9+, three samples were additionally ARX+. The SOX9- Group B sample was additionally ARX- and PDX1- (Fig. S4f). Our IHC analyses showed that α-/β- like tumors in Group A harbored significantly more ARX+PDX1-SOX9-, while Group B acinar-like tumors were enriched for ARX-PDX1-SOX9+ features (adjusted p-value = 0.001998: Fisher’s Exact Test and fdr corrected).

## Discussion

Here we report a methylation-based classification which accurately distinguishes PanNECs from all PanNENs, including G3 PanNETs. Our study demonstrates a potential exocrine cell of origin of PanNECs, distinct from the endocrine cell of origin of PanNETs. Our findings lead to a novel approach extending the classical histopathological diagnosis by an epigenomic diagnosis of two otherwise histologically challenging aggressive subsets of PanNEN.

DNA methylation at CpG dinucleotides is a mechanism of cell-type specific gene regulation inherited in a continuous manner throughout development, hence it is a robust marker of cell identity ^50^. Methylation patterns are now considered as a robust method to identify tumor cell-of-origins across tissues ^51^, and in different cancer types ^47,52–55^ . In addition, they have been used for characterization of subgroups within a tumor entity, for example in central nervous system tumors ^56^, breast cancer ^53^, ovarian cancer ^57^ as well as tissue origin of cell-free DNA (cfDNA) in disease ^48^. Furthermore, methylation analysis has shown the significance of hypermethylation during the transition of committed cells to a naive stem cell state ^44^. The cell type from which a cancer originates is highly informative to its identification, classification, treatment and prognosis ^58^. To this end, we utilized methylation profiles of PanNEN tumors in order to demonstrate cell-of-origin identity.

Our investigation of 10K most variable probes clustered a PanNEN cohort, representing all tumor grades, into three groups. The largest two groups, Groups A and B, clearly distinguished NECs from the rest of the cohort. Three recent studies have investigated PanNET subgroups at the epigenetic level ^17–19^. Cejas et al. looked at histone acetylation and transcriptomes ^17^ of NET samples, with two neuroendocrine samples both from the ileum. Di Domenico et al. ^18^ and Lakis et al. ^19^ used 450K methylation profiles from cohorts of well-differentiated PanNET samples. Our work expands the field by including the most aggressive subtypes of PanNENs: NETG3 and NEC tumors, which the aforementioned studies largely excluded (Di Domecio et al. included only two NETG3). In addition, we used the 850K EPIC beadchip assay, significantly increasing the number of CpGs investigated. All three studies identified the cell of origin of early PanNETs to α and β cells of the pancreas. These findings are reflected in our analysis of tumors belonging to Group A. Due to the inclusion of more high-grade NETG3 and poorly differentiated high-grade PanNEC samples in our study, the methylation patterns separate the groups into more encompassing subgroups, placing those of low grade, and α and β similarity together, distinct from the undifferentiated PanNECs (Fig.1d).

Phylo-epigenetic analysis exposed the tight clustering of PanNEC samples with exocrine cells and their separation from endocrine cells and their relationship to PanNET tumors (Fig. 4a). PanNECs have been repeatedly compared to PDACs in terms of mutational spectrum ^14^. The key difference became obvious at the cell-type similarity level whereby PDACs were clearly similar to ductal and acinar cells, whereas NEC profiles were only similar to acinar cell profiles (Fig. 4c). A recent study by Kopp et al. showed that SOX9 accelerated the formation of precursor lesions of PDAC when co-expressed with oncogenic KRAS. By lineage tracing, their study also suggested that upon the expression of *SOX9*, Pancreatic Intraepithelial Neoplasia (PanIN) lesions and subsequently pancreatic ductal adenocarcinoma arise from ductal metaplasia of the pancreatic acinar cells, a phenomenon known as acinar-to-ductal metaplasia ^42^. SOX9 is a crucial factor regulating pancreatic cell development, initially maintained in the multipotent progenitor state ^25,29^, and subsequently restricted to NKX6.1+ bipotent progenitor cells. Later, in adult pancreatic cell types, SOX9 is constrained to ductal and centro-acinar cells of the pancreas ^27,40^. We detected SOX9 in 81% (9 /11) of PanNECs available for IHC in Group B and 60% (3/5) of NETG3 samples in Group A. Interestingly, Group B samples carried a SOX9+, ARX- and PDX1- phenotype (Fig. S4f) and mirrored the naive hESC methylation profile (Fig. 1g). In line with the aforementioned findings in PDAC, our data led us to hypothesize that Group B samples are acinar-like tumors which undergo a mechanism similar to that of PDAC formation via the expression of *SOX9*. In contrast, NETG3 SOX9+ samples found in Group A carried a profile similar to α cells but not to acinar cells (Fig. S5c). Since SOX9 plays a critical role in the multipotent and bipotent state of pancreatic development, these NETG3 tumors may originate from endocrine cells, given their similarity to α and β cells and in the course of tumor progression, revert to expressing SOX9 as a mechanism to move towards a progenitor-like phenotype.

PanNEC mutational patterns such as *KRAS*, *SMAD4* and *TP53* were present in Group B. Using focal DNA copy number analysis we found that Rb1 loss, shown in NEC tumors at the protein level ^59^, was due to DNA copy number loss of chromosome 13, found in 50% of the PanNECs but not in any NETG3 samples (Fig. 2f). PanNEC subgroups with KRAS and Rb1 loss showed a higher response rate to first line platinum-based treatment, but with a shorter overall survival rate ^60^. In contrast, cell cycle regulators *CDKN1B and CDKN2A* were lost in Group A PanNET samples. These important driver events in NETs and NECs indicate a conversion of both tumor entities to cell cycle regulation pathways and could indicate separate directions of drug treatment (reviewed by Scarpa ^61^).

Group A tumors were all PanNETs, and included all grades G1, G2 and G3 with mutated *ATRX*, *DAXX* and *MEN1* (A-D-M) genotypes as well as known copy number patterns for PanNET tumors^62^. The top DMPs in pancreatic cell markers belonged to *IRX2* and *NKX6-1*, and maintained a strict hypomethylation in Group A, strongly indicative of an endocrine origin (Fig. 3b). IRX2 and PDX1 characterized α-like and β-like tumors, respectively ^28–30^, and a closer look at their methylation pattern in Group A clearly separated the samples into an α-like subgroup (*PDX1* hypermethylation and *IRX2* hypomethylation), and a β-like and intermediate subgroup of endocrine-like tumors (*PDX1* hypomethylation and *IRX2* hypomethylation) that carry >75% β cell signature or equal proportion of α, β cell signature, respectively (Fig. 3c; Fig. S5d). The mutational characteristics of α-like tumors identified by Chan et al. is reflected in our results, where the majority of samples with A-D-M mutations show promoter hypermethylation in *PDX1* and hypomethylation in the *IRX2* gene probes and strong alpha-cell type signature ^45^. (Fig. 3c; Fig. S5d). In addition, the α-like subgroup showed ARX+ PDX1- expression in 12/13 stained samples, further validating their cell-of-origin identity. The β cell-like and intermediate-like tumors however failed to display a consistent IHC expression pattern of ARX and PDX1. The β-like cell signature tumors were in fact insulinomas (samples PNET 25, 52, 91) that showed an ARX+ PDX1- profile, further highlighting that methylation is a strong marker of lineage, and more likely to expose cell of origin than expression of these markers. Nevertheless, the question remains as to how insulin is maintained in the absence of PDX1, a major transcription factor controlling insulin production in β cells ^63^. The intermediate tumors included PDX1 and ARX double negative (PNET 51, 50 and 18), PDX1 positive only (PNET 51) and ARX positive only (PNET 14) (Fig. S4d).

Group C tumors, with one NETG2, one NETG3 and two PanNECs, had no mutations picked up by our targeted panels (Fig. S2b) and had a methylation pattern of CpGs associated with cell markers resembling a mixed Group A and B profile (Fig. 3b). Although they remain close to Group B samples in the phylo-epigenetic analysis (Fig. 4a), their similarity to acinar cell signatures is not comparable to that found in Group B (Fig. 4c). Further information from Group C is limited due to the small number of samples.

Our work establishes an exocrine cell of origin for PanNECs resembling an acinar cell type. Methylation profiling is a superior method of tumor-type identification to genomic mutations, copy number alterations, or IHC of single markers. This is due to the epigenetic memory of the cancer’s cell of origin, as discovered in several studies ^64,65^, most importantly in Cancers of Unknown Primary (CUP) ^48,55^. Epigenetic identification of cancer cell-of-origin can determine the diagnosis ^18,56^, evolution ^66–68^ and treatment ^60,69^ of this disease. Our study supports the introduction of methylation analysis in routine diagnosis of high-grade PanNEN tumors.

## Materials and Methods

### Patient Cohort and experimental representation

Our cohort consists of 57 PanNEN samples collected from 55 patients. For one patient (PNET77), both primary and metastasis was obtained and for another patient (PNET56), two metastases were obtained from two different time points, one year apart. The Institute of Pathology at Charité - Universitätsmedizin Berlin provided 48 samples of all grades and the University of Bern provided 9 NETG3/NEC samples. We also had one metastasis each in the bladder, papillary and peritoneal regions. The primary tumors resected were located in the head (29.8%), tail (24.6%) and body (3.5%) of the pancreas. Two samples had lesions in multiple sections of the pancreas. Clinical reports on the tumors were collected and are presented in Supplementary table 1. All samples were collected as Formalin-fixed paraffin-embedded (FFPE) blocks, and normal controls for the respective patients were collected with the exception of 10 cases (Fig. 1a). Normal tissue sections were obtained as either tissue adjacent to the tumor (as per the pathologist’s examination) (normal adjacent n= 15, Fig. S1b), or as a completely separate block containing only normal tissue (normal distant n = 29). Normal blood samples were available in 3 cases. All patients provided signed consent as part of the clinical documentation protocol of the Charité - Universitätsmedizin Berlin. Samples of the University of Bern were provided by the Tissue Biobank Bern (TBB) according to the relevant Ethics approvals.

### PanNEN panel design

We designed a custom panel targeting PanNEN relevant genes, covering all mutations associated with PanNENs. The designing involved first performing a text-mining approach to extract high-quality information regarding genes associated with PanNENs from GeneView^48^, the Catalogue of Somatic Mutations in Cancer (COSMIC)^49^ and mutations collected from PanNEN publications ^14,50–52^. From this list, 47 PanNEN-likely driver genes were then extracted with further focus on MAPK and mTOR pathways. Amplicons were then designed for the GRCh37 genome by providing the Ion AmpliSeq Designer tool (Life Technologies) the candidate genes under the criteria “DNA Gene design (multi-pool)” . The panel was designed to generate primers targeting 125-bp stretches of exon regions of the selected genes. The complete panel included 1175 amplicons, divided into two pools.

### DNA isolation

DNA from all samples was isolated from tissue that had been fixed as FFPE. Tissue samples were sectioned and stained with Hematoxylin and Eosin (H&E). Pathologists demarcated tumor and healthy tissue areas in the H&E slides and depending on the size of the marked area, 12 sections of 5µm each from tumor samples and 6 sections of 5µm for control normal tissue were used for DNA isolation (Fig. S1b). The tissue was macro-dissected from the slides and DNA was prepared using the GeneRead DNA FFPE kit (Qiagen, Netherlands). Quality and quantity of DNA was determined by RNAse P quantification (Thermo Fisher Scientific, USA).

### DNA Sequencing

We used 20ng of DNA for library preparation using Ion Ampliseq Library kit (Thermo Fisher Scientific). Regions were targeted by primers distributed into two amplicon pools per DNA sample for PanNEN panel and 4 amplicon pools per DNA sample for Comprehensive Cancer Panel (CCP). Upon ligation to Ion Xpress Barcode Adapters (Thermo Fisher Scientific) and purification using Agencourt AMPure beads (Beckman Coulter), two samples were mixed at equal ratio on a 318v2 sequencing chip. Using the Ion Torrent PGM (Thermo Fisher Scientific), the samples were sequenced at an average read depth of 1158 reads using PanNEN panel and an average read depth of 217.03 reads for CCP, in order to generate the raw intensity data.

### Sanger Sequencing

To validate results from targeted massive parallel sequencing we performed Sanger sequencing on specific mutations which we found required validation, for example sub-optimal amplicon performance or location within long nucleotide repeat areas. The resulting signal intensity images were manually scanned to identify the targeted mutations (Supplementary table 4).

### Fluorescence in-situ hybridization (FISH)

Fluorescence in situ hybridization (FISH) was performed on 3μm tumor sections from 23 samples. We used commercially available, standardized probes for detecting chromosome 5 with a *RICTOR* gene probe, chromosome 9 with *TGFBR1* gene, and chromosome 11 with *MEN1* gene (Empire Genomics, USA). Hybridization was performed according to manufacturer’s instructions. Where possible, we scored 40 cells per sample using an Olympus microscope. Analysis was conducted using ‘BioView solo’ (Abbott Molecular).

### DNA methylation

DNA methylation profiling of all PanNEN cohort samples was performed with 200–500ng of DNA using the Infinium® MethylationEPIC BeadChip array (850K; Illumina, Inc., San Diego, CA, USA) according to the protocols provided by the manufacturer.

Normal cell type 450K methylation data of Pancreatic α, β, acinar and ductal cells were obtained directly from the published lab or GEO (Neiman et al, GSE122126 and GSE134217). In addition, published 167 PDAC 450K methylation data was also obtained from GSE49149. We also obtained hESC EPIC data from GSE128130. All datasets utilized are shown in the table below.

| Sample ID | New sample annotation | Dataset |
| --- | --- | --- |
| Alpha rep 1 | Alpha A | Neiman et al. |
| Alpha rep 2 | Alpha B | Neiman et al. |
| Beta rep 1 | Beta A | GSE122126 |
| Beta rep 2 | Beta B | GSE122126 |
| Beta rep 3 | Beta C | Neiman et al. |
| Ductal 1 | Ductal A | GSE134217 |
| Ductal 2 | Ductal B | GSE134217 |
| Ductal rep 1 | Ductal C | GSE122126 |
| Acinar 1 | Acinar A | GSE134217 |
| Acinar 2 | Acinar B | GSE134217 |
| Acinar rep 1 | Acinar C | GSE122126 |

### Data processing and analysis

#### Mutational analysis

The raw reads were first aligned to GRCh37 sequence using the Torrent Mapping and Alignment Program (TMAP; Life Technologies), placing a cut-off of > 50 nucleotides in aligned reads and a mapping quality of > 4 using an in-house python script. The processed bam files were then utilized for variant calling using TS Variant Caller plugin under the “strict” setting per Ion Suite (Ion Torrent platform) parameter profiles. The generated variant call format (VCF) files of the tumor-normal pair per patient was merged and reference and alteration read count for all variants within the two VCF files were extracted to determine the representation of the variants in both cases. The merged VCF file was then annotated using the SoFIA^53^ annotation framework. Downstream filtering included removing variants that were positive for the following set of parameters: intronic and synonymous variants, 1000Genome variants with frequency greater than 1% in the population, variants within homopolymer regions > 4 nucleotides, tumor variant allelic frequency < 5% and finally, variants that were represented at comparable allelic frequencies in the matched normal tissue.

#### DNA methylation preprocessing

Raw idat files were preprocessed using subset-quantile within array normalization (SWAN) provided through the R package minfi^54,55^. Probes performing poorly in the analysis were further filtered out. The failed probes were identified if their detection p value was > 0.01 in at least one sample. Probes cross reactive to multiple sites in the genome^56^, Sex chromosome probes and probes containing SNPs with an allele frequency > 0.01 were also filtered out.

When comparing 450K and 850K samples, the processing was done by first converting the 850K platform to 450K and then performing the aforementioned steps. When tumor and normal data were analyzed together, normalization was done using the preprocessFunnorm() function from the minfi package in order to accommodate for the global variation between normal and tumor data. This also entailed removing cross reactive probes of EPIC and 450K array (https://github.com/sirselim/illumina450k_filtering). hESC data normalization was performed as depicted in Patani, *et al,* 2020. Briefly, we normalized the data using the preprocessNoob() function from minfi and performed the aforementioned filtering approach to remove problematic probes. Finally, beta values for each dataset were used for all downstream analysis, statistics and visualization.

#### Subgroup identification and associated analysis

Using the 10K most variable probes identified by determining row (probe) standard deviation (σ), DNA methylation-based classes of PanNEN were identified with the R package ConsensusClusterPlus under the following parameters: maxK=12, reps=1000, pItem=0.8 and pFeature=1. The function performed agglomerative hierarchical clustering after performing 1-Pearson correlation distance. A consensus matrix carrying pairwise consensus values was finally generated for 12 clusters. The most stable number of clusters was determined based on the cumulative distribution score curve (CDF) that reached approximate maximum (k=3) and the correlation heatmap for each 3-mer. Hierarchical clustering of 10K variable probes was done by first obtaining a dissimilarity matrix using an Euclidean algorithm and then performing the clustering using complete linkage.. Hierarchical clustering was done using the R package “Stats” and tSNE was done using the “Rtsne” package under the perplexity=8. Genes associated with the 10K most variable probes were evaluated for GO pathway ^57^ biological processes term enrichment using the enrichGO() function in the clusterprofiler R package. The analysis was done under the following parameters: pAdjustMethod = “BH” (Benjamini and Hochberg), pvalueCutoff = 0.01, qvalueCutoff = 0.05. All genes represented in the Illumina EPIC array were used as background. In order to reduce generality of GO terms, the simplify() function was used. The final set of terms was curated by filtering only those that showed an adjusted p-value less than 0.05 and fold enrichment of greater than 1.5. -Log_10_P value was calculated for the remaining terms and a barplot was generated for the 12 most significant terms using the ggplot2 package.

#### Differentially methylated probes (DMP) and associated analysis

Upon extracting and assigning the samples to the identified stable clusters Group A and Group B, differentially methylated probes (DMP) were identified out of all the CpG sites (upon the aforementioned preprocessing) using CHAMP package function champ.DMP() under the following parameters: adjPVal = 0.05, and adjust.method = “BH”, arraytype=”EPIC”.

Differentiated pancreatic cell markers were curated from PangaloDB (https://panglaodb.se/). Cell markers showing sensitivity_human > 0.05 for α, β, Ɣ, δ, Epsilon, Acinar, Ductal and Islet Schwann cells were obtained. DMP associated genes that overlapped with curated Islet cell markers were extracted. Final lists of DMP associated Islet cell markers were identified if they met the following criteria: abs(Δbeta) > 0.25 and -log_10_P > 5. To determine closely related samples within each group, hierarchical clustering with complete linkage was performed using beta values of the identified Islet cell markers associated with DMPs before visualization.

DMP between α, β, ductal and acinar cell types were identified using CHAMP package function champ.DMP under the following parameters: adjPVal = 0.05, and adjust.method = “BH”, arraytype=”450K”. Significant probes of each DMP set showing absolute Δbeta value > 0.2 and adjusted p-value < 0.01 were obtained. A total of 46,500 unique probe IDs were collected and defined as DMPs of normal cell types. Upon preprocessing and downstream filtering (see DNA methylation preprocessing section) of tumor and normal data combined, 38892 DMPs of normal cell type overlapping in tumor-normal matrix remained and methylation values were extracted to calculate Pearson distance using the function get_dist() from factoExtra R package. Finally, neighbor-joining tree estimation was performed using nj() function in the ape package to generate phylo-epigenetic trees. hESC probe identification was performed using an adaptation of method in Patani, *et al.,* 2020. Briefly, hESC probes carrying a mean beta < 0.3 across the primed cells were retained as background matrix. Unmethylated probes of hESC were then defined as those that carry mean beta < 0.3 in both primed and naïve hESC. Hypermethylated probes of hESC were defined by first calling DMPs using the background matrix. CHAMP.DMP() ran with the adjPval=0.05, adjust.method=“BH”, and arraytype=“EPIC”. Probes carrying Δbeta < -0.1 (hypermethylation in naïve state compared to primed state) were then extracted. Finally, to compare to the tumor samples, after normalization, preprocessing and filtering (see above section: DNA methylation preprocessing), the methylation values for unmethylated probes of hESC and hypermethylated probes of hESC in the tumors were determined and the mean per probe type in each sample was computed. The distribution of these computed mean per Group was visualized. In addition, mean values for NETG3 and NEC samples of Group A and Group B were extracted and separately visualized.

#### Normal cell signature analysis

In order to determine cell signature proportion in each sample, the methodology provided by Moss et al. was utilized. Briefly, first the reference atlas (https://github.com/nloyfer/meth_atlas) was obtained, then just the β, ductal and acinar cell profiles and their featured CpGs were extracted. In order to add an α-cell profile, the normal cell types were preprocessed and normalized (Neiman et al., GSE122126 and GSE134217), and the α cells were extracted. The mean value of each probe was calculated and the overlap of the probes compared to the featured CpGs of Moss et al. were obtained. The final matrix of the normal reference “atlas” contained the methylation values for α, β, ductal and acinar cells of the overlapping probes. The euclidean distance between each sample given the probes was computed using get_dist() function from the FactoExtra R package. PanNEN and PDAC data were normalized separately as mentioned above, and the methylation beta value matrix was transformed for subsequent analysis. The program developed by Moss et al. (https://github.com/nloyfer/meth_atlas/blob/master/deconvolve.py) was then employed to identify the normal cell signature proportion in samples of the PanNEN and PDAC cohorts.

#### Copy Number Aberrations (CNA)

CNA was identified from EPIC array data using the R package conumee^58^. Upon raw preprocessing, mean of similar CNA segments per autosomal region were obtained and a mean value per autosome was calculated for each sample to determine the log2 ratio of intensities across the chromosome. A cut-off of x > 0.15 and x < -0.15 was placed to limit the number of false positives obtained upon comparing log2 ratio values to FISH count (Fig. 2d). To determine the subgroups within the cohort, euclidean hierarchical clustering was also performed on the data. To determine focal aberrations, the chromosomal segment log2 values, determined by conumee were obtained for samples of each group and were separately run in GISTIC software under the following parameters: -genegistic 1, -smallmem 1, -broad 1, -brlen 0.5, -conf 0.90 -armpeel 1 and -gcm extreme. GISTIC was only performed in Group A and Group B, and not for Group C, due to the limited number of samples.

### Immunohistochemistry (IHC)

Representative samples from Group A and B were subjected to Immunohistochemistry (IHC) for ARX, PDX1 and SOX9 expressions. 2.5µm FFPE sections were used for ARX (1:1500, R&D Systems, sheep, AF7068), PDX1 (1:100, R&D Systems, mouse, MAB2419)) and SOX9 (1:100, Cell Signaling, rabbit, mAb #82630) immunostainings. Antigen retrieval was performed by heating the Tris30 buffer at 95°C for 30 minutes. The primary antibody was incubated for 30 minutes at the specified dilutions. Visualization was performed using Bond Polymer Refine Detection kit, using DAB as chromogen (3,3’-Diaminobenzidine). In most of the cases staining was identified as strong positivity for the specific TFs, single cell positivity for few cases was assessed and is reported in Supplementary table 1. The immunostaining for all antigens was performed on an automated staining system (Leica Bond RX; Leica Biosystems, Nunningen, Switzerland).

### Data visualization and statistics

All data analysis, statistics and visualization was performed in R (version 4.0.0). Visualization was done using the base R plotting function, ggplot2 package or ComplexHeatmap package. The appropriate statistics mentioned above were all performed using respective R packages or base R functions. For survival analysis, the “survival” and “survminer” packages were used ^70,71^.

## Acknowledgments

We would like to acknowledge Peter Mohr for work on the NGS panel. Torsten Gross, Thomas Sell and Manuela Benary for their help with support in coding and data analysis. Renaud Maire, Daniel Teichmann, Kerstin Wanke-Möhr, Manuela Pacyna-Gengelbach, and Hatice Noyan for technical assistance. Philip Bischoff for pathology support. Jeroen van Marle for language editing.

## Contributions

TS: study concept and design; analysis and interpretation of data; statistical analysis; technical or material support; writing of the manuscript. PR: critical revision of the manuscript for important intellectual content; writing of the manuscript. ADD: IHC and analysis. FB: interpretation of data and methylation analysis pipeline; critical revision of the manuscript for important intellectual content. AM: technical and material support. SK: PanNEN DNA sequencing panel generation. DL: analysis and interpretation of FISH data. LHC and UL: PanNEN DNA sequencing panel generation. KD: clinical analysis. FR and AJ: clinical analysis. MM: critical revision of the manuscript for important intellectual content. NB: interpretation of data and study supervision. MP: clinical analysis and interpretation of data. DH: analysis and interpretation of clinical data. DC: analysis and interpretation of data, technical support. IM: sample collection, analysis and interpretation of data. AP: sample collection, analysis and interpretation of data, critical revision of the manuscript for important intellectual content, SM: study concept and design, writing of the manuscript; study supervision; funding. CS: study concept and design; critical revision of the manuscript for important intellectual content; study supervision; funding.

## Ethics Statements

Tissue collection obtained from Charité Universitätsmedizin Berlin was processed according to the Charité ethics vote EA4/022/15. PanNEC samples from Bern with ethics approval: KEK_BE 105/2015.

---

**Fig. S1.**
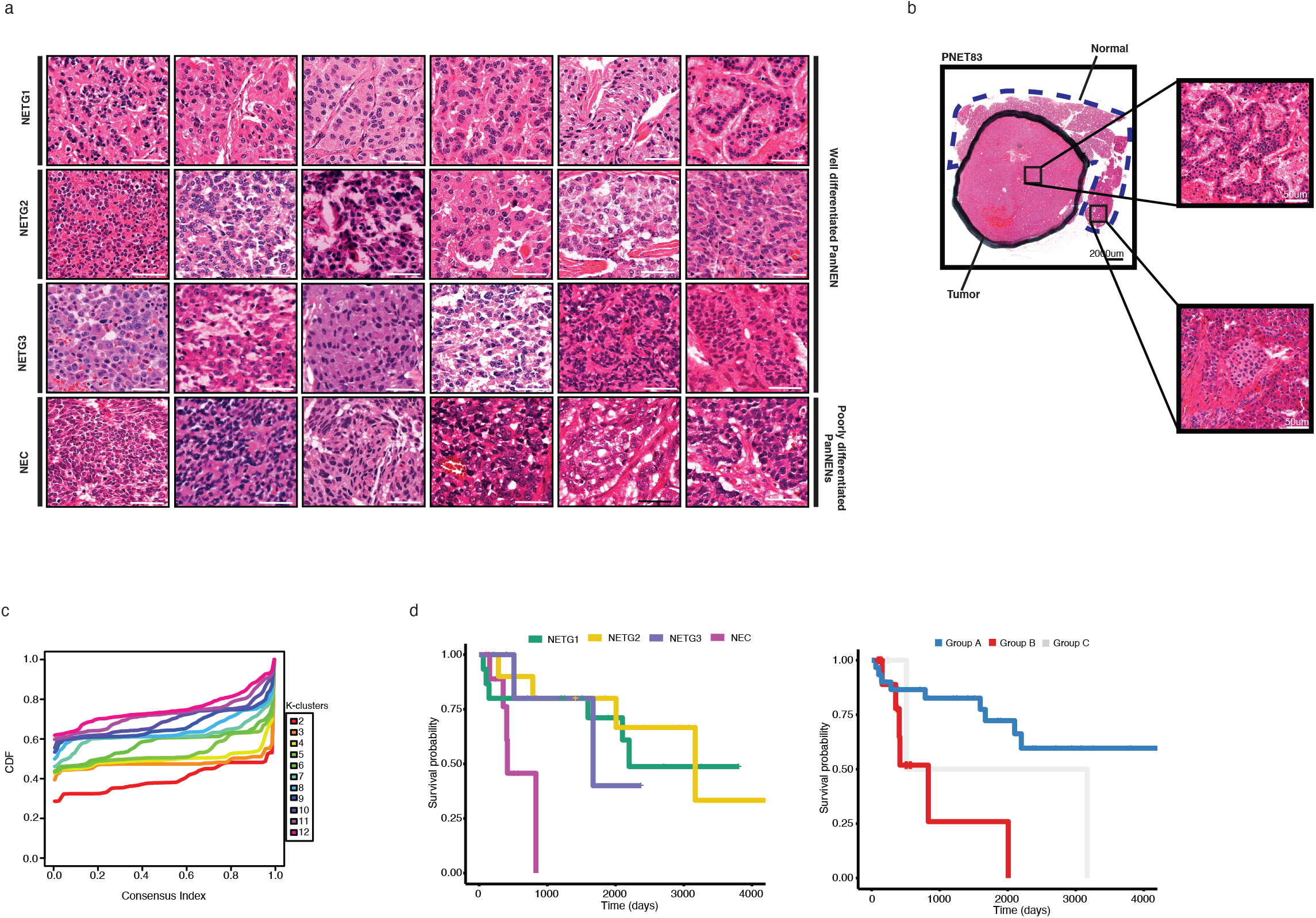
Three main subgroups define the PanNEN cohort. **a.** Representative H&E stainings for each grade. **b.** Representative macro images of tissue H&E sections used for downstream analysis. Solid and dotted lines represent tumor and normal regions respectively, as per a pathologist’s evaluation. **d.** Cumulative distribution function (CDF) curve of the resulting ‘k-mer’ count for 12-k’s. Cluster count of ‘three’ resulted in the most stable number subgroups. Scale bar: 50µm. **c**. Kaplan Meier survival graph from available patient data. Left: graph shows cohort according to tumor grade (P = 0.062). Right: graph shows cohort according to methylation groups A, B and C. Significant difference found between Group A and B (p = 0.0049) and between all groups (p = 0.011). P value generated using log-rank sum test statistics by survminer R package.

**Fig. S2.**
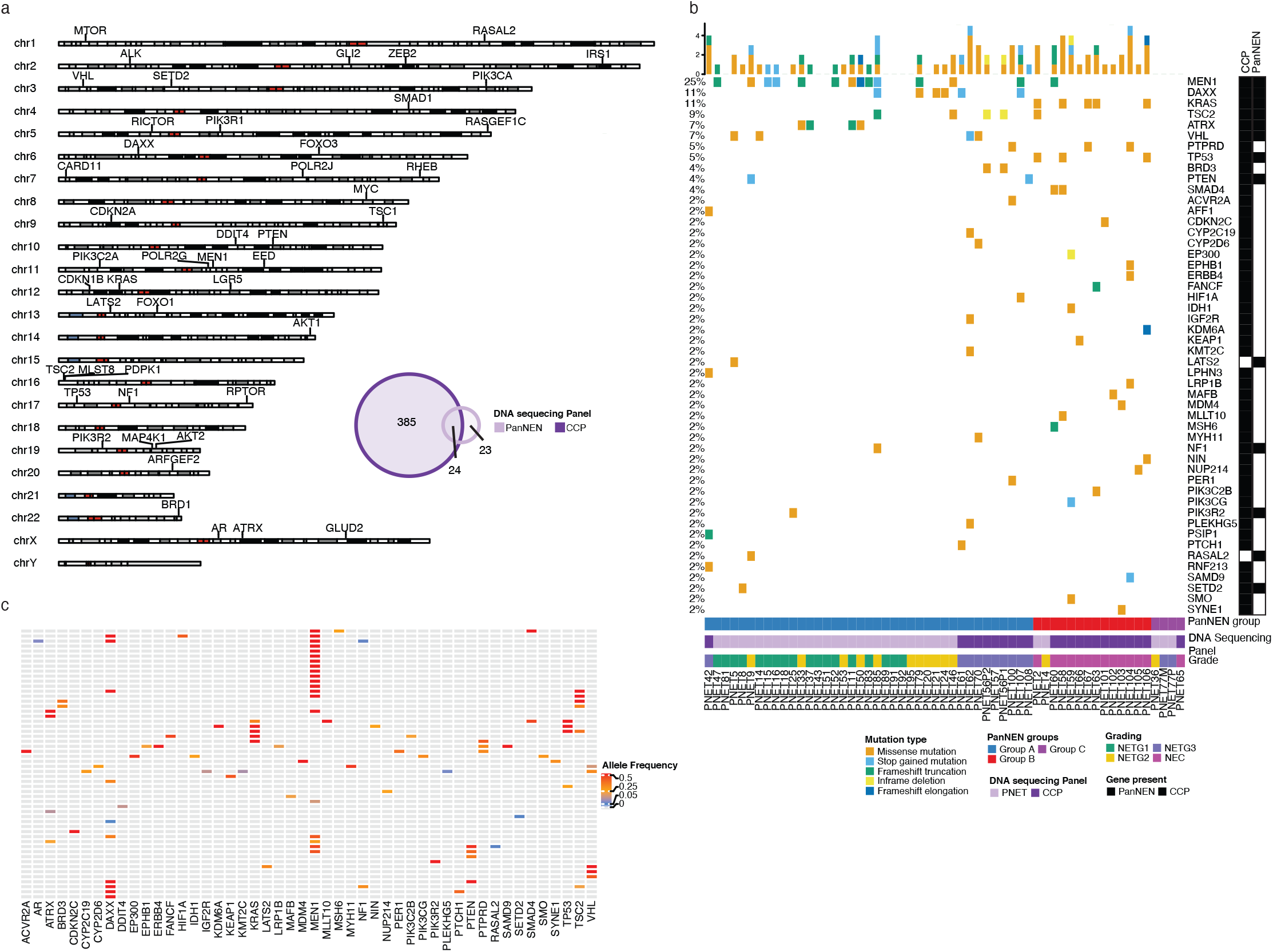
Distinct mutational aberrations define PanNEN subgroups. **a.** Custom PanNEN gene panel for mutation analysis. Targeted genes of PanNEN panel are displayed in their respective chromosomal region. PanNEN panel and commercial Comprehensive Cancer Panel (CCP). **b**. Mutational profile of PanNEN cohort. (top panel) Barplot depicting mutational frequency in a given sample, colored according to variant type; (left) Percentage value per row: frequency at which the respective gene is aberrated in the cohort; (middle panel) mutation variant of each patient, colored according to variant type; White spacing: no mutation was identified in the regions of the targeted gene; (right) two row annotations displaying whether the respective gene was covered by PanNEN panel, CCP panel or both; PanNEN subgroup, tumor grade and gene panel used for mutational analysis for each sample is annotated at the bottom. **c**. Allelic frequency of the mutations in the PanNEN cohort. Mutated genes are displayed at the bottom; Heatmap color represents the frequency at which the alterations were identified in a given sample (range from: 0.05 [blue], to 0.93 [red]). Grey: absence of aberration in the respective gene for a given sample.

**Fig. S3.**
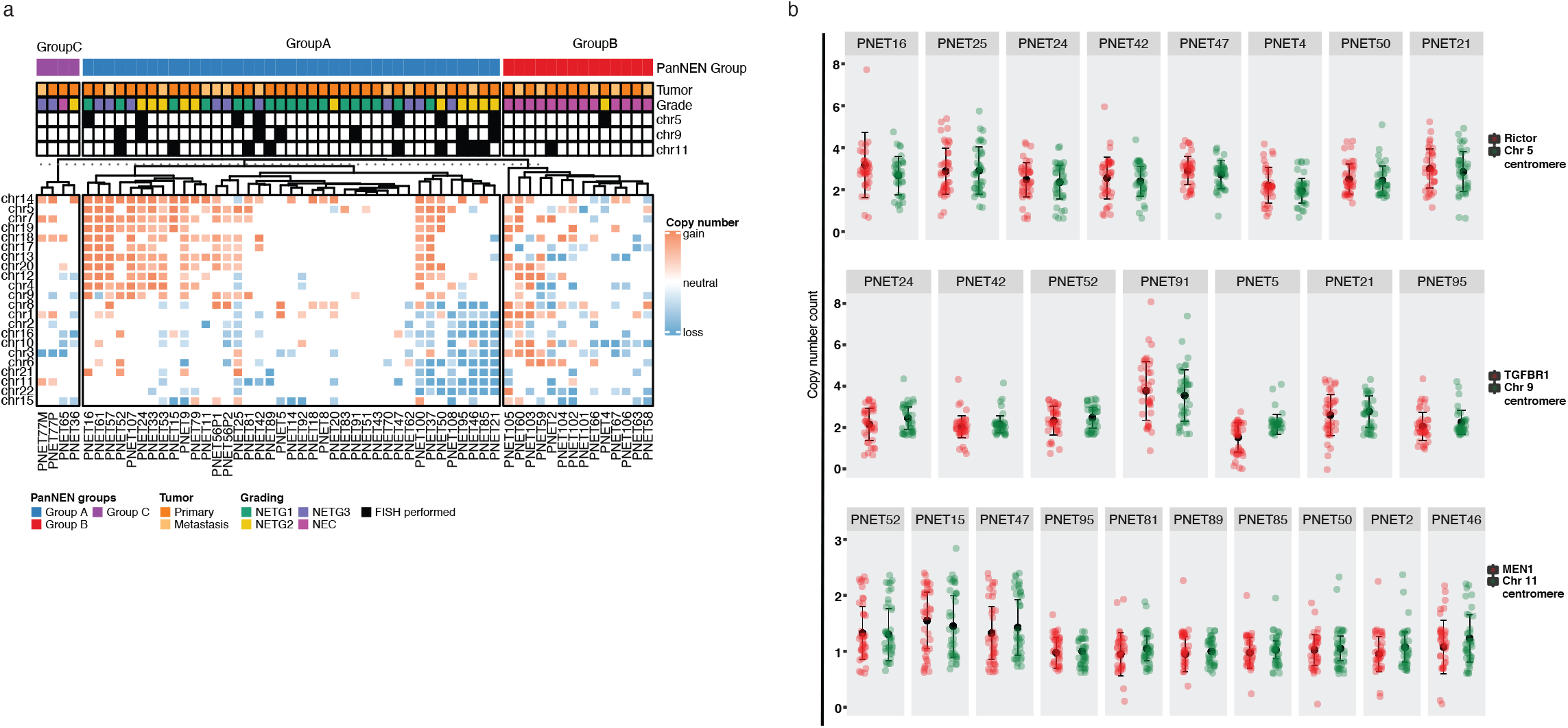
Whole chromosomal aberrations distinguish PanNEN subgroups. **a.** Hierarchical clustering of mean log2 ratios of chromosomal segments per autosome; column annotation (bottom): tumor grade, tumor type in addition to FISH validation status; representative samples were analyzed for gains in chromosome 5 and 9 and loss in chromosome 11 (black boxes in the respective column annotations). **b**. Quantification of chromosomal aberrations for chromosomes 5, 9 and 11 was performed using (red) by FISH. n = 40 cells per sample; The distribution of signals per sample is depicted; Each point represents the mean count of cells for the respective probe; Black: mean, error bars standard deviation.

**Fig. S4.**
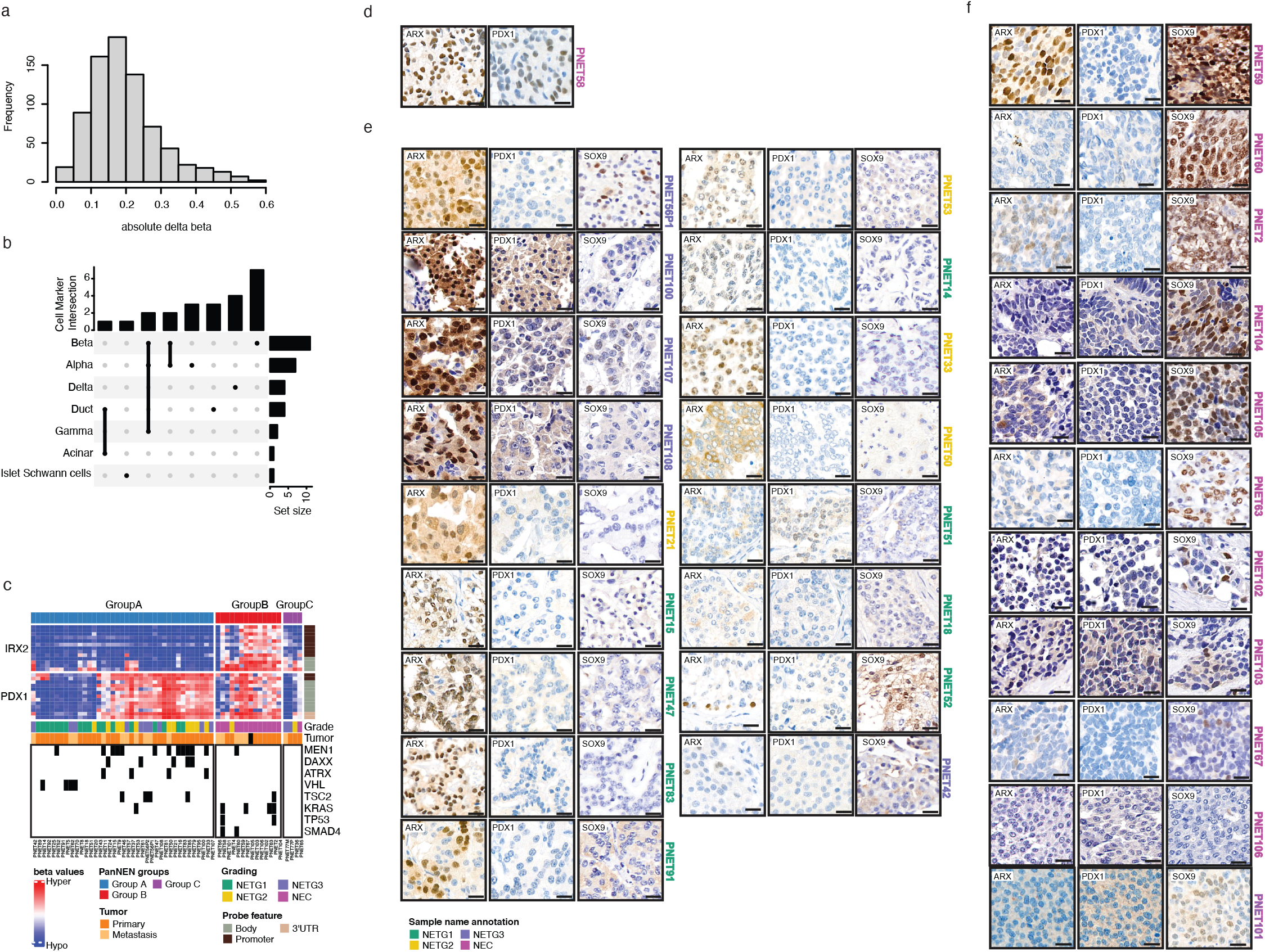
Pancreatic cell markers define PanNEN subgroups. **a.** Distribution of absolute log2FC of pancreatic cell marker associated DMPs. Absolute log2FC ranges from 0.03 to 0.59 (x-axis), frequency of a given absolute log2FC on y-axis. **b.** Number of markers associated with each cell type represented in the bar plot; Markers associated with multiple cell types are also depicted; the cell type of the respective associations are linked (bottom annotation). **c.** IHC of ARX, PDX1 and SOX9 in Group A. All samples that underwent IHC for all three markers are shown. **d**. IHC analysis of ARX, PDX1 and SOX9 in Group B. The samples that underwent IHC for all three markers are shown. Scale bar: 20µm.

**Fig. S5.**
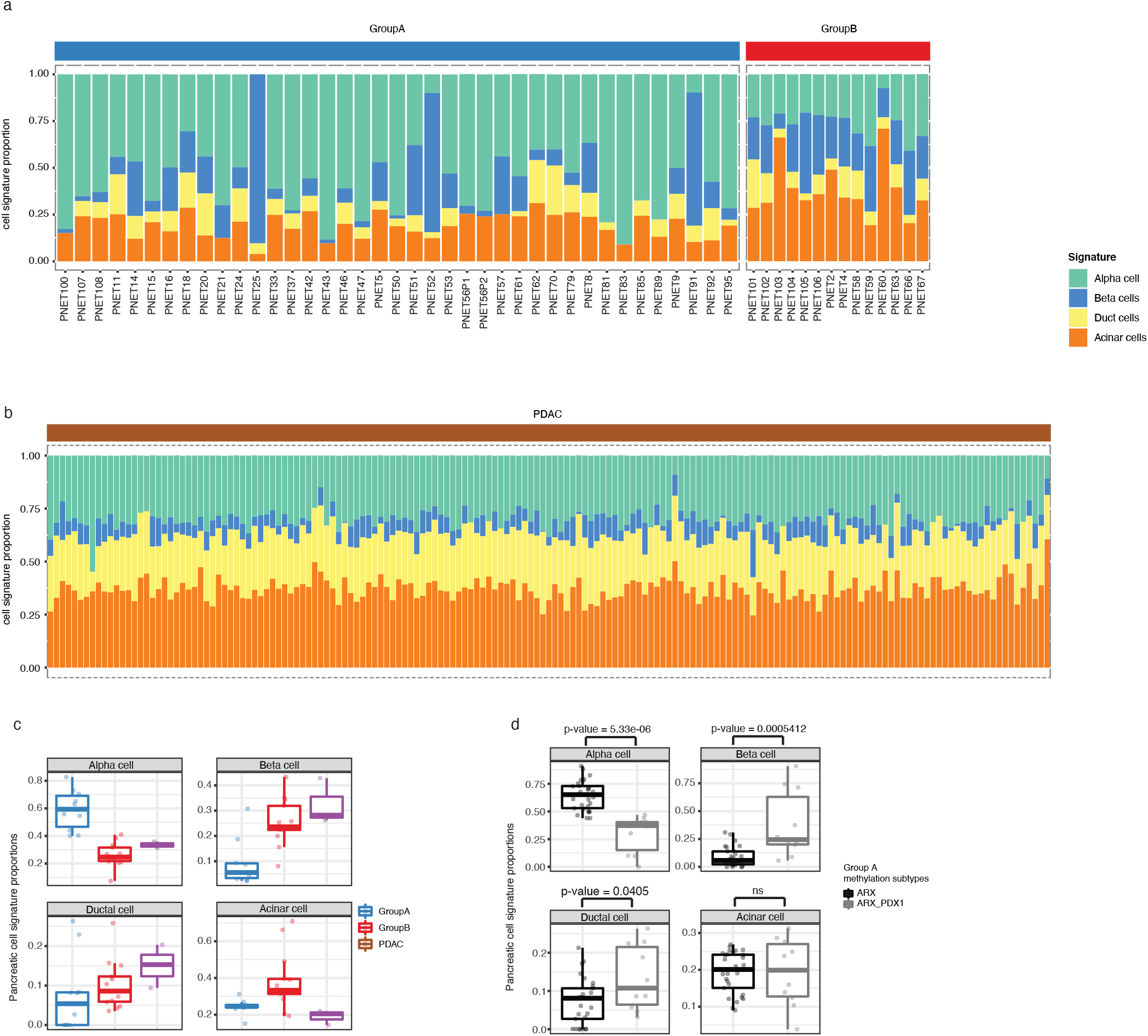
Pancreatic cell atlas signatures define PanNEN subgroups. **a.** Stacked bar plot representing proportions of the pancreatic cell signatures; top annotation: PanNEN subgroup for the respective samples. **b.** Proportion of pancreatic cell signature in each PDAC sample (n=167). Legend as in a. **c**. Proportion of pancreatic cell atlas signatures in NETG3 and NECs of Group A and Group B, as well as PDACs. Boxplot representing distribution of the proportion of atlas signature of α-, β-, ductal and acinar cells (each main box) in the NETG3 and NECs of subgroups and PDACs; Each dot depicts the proportion of atlas signature of the respective cell type in a given sample. **d**. Proportion of pancreatic cell atlas signatures in *IRX2* hypo/*PDX1* hypermethylated and *IRX2* hypo/*PDX1* hypomethylated tumors of Group A. Boxplot represents distribution of the proportion of atlas signature of α-, β-, ductal and acinar cells (each main box) Group A tumors of respective *ARX* and *PDX1* methylation profile; Each dot depicts the proportion of atlas signature of the respective cell type in a given sample.

